# A direct estrogenic involvement in the expression of human hypocretin

**DOI:** 10.1101/2023.12.21.572761

**Authors:** Haimei Li, Xinlu Chen, Jingyi Dong, Ripeng Liu, Jinfeng Duan, Manli Huang, Shaohua Hu, Jing Lu

**Affiliations:** Department of Psychiatry, the First Affiliated Hospital, Zhejiang University School of Medicine, Hangzhou, Zhejiang 310003, China; The Key Laboratory of Mental Disorder Management in Zhejiang Province, Hangzhou, Zhejiang 310003, China; Department of Psychiatry, Sir Run Run Shaw Hospital, Zhejiang University School of Medicine, East Qingchun Road 3#, Hangzhou, Zhejiang 310016, China; College of Life Science, Zhejiang Chinese Medical University, Hangzhou, China; College of First Clinical College, Zhejiang Chinese Medical University, 548 Binwen Road, Binjiang District, Hangzhou 310053, China

**Keywords:** Estrogen receptors, hypocretin, sex difference, depression

## Abstract

**Background:** Hypocretin-1 may play an important role in depression, which was increased in female depression in LHA in our previous study. Estrogen acts through its nuclear transcriptional receptors.

**Methods:** We studied the possibility of a direct action of estrogen receptors on the expression of human hypocretin. To explore the activation of ERs by hypocretin, we first investigated the potential presence of co-localization of the estrogen receptors (ERs) in hypocretin-immunoreactive neurons in the lateral hypothalamic area (LHA) in human and nuclear ERs in hypocretin immunoreactive neuroblastoma SK-N-SH cells. We investigated the relationship between E2 and hypocretin in female rats. After we explored the regulating role of exogenous estrogen on hypocretin gene expression via ERs through Chromatin immunoprecipitation, Electrophoretic mobility shift assay and Luciferase assay.

**Results:** We found that hypocretin-1 plasma level was significantly higher in female depression. Estrogen receptors (ERα and ERβ) and hypocretin are co-localized in female depression LHA, PC12 and SK-N-SH cell lines. The estrogen receptor response elements (EREs) existing in the hypocretin promoter region may directly regulate the expression of hypocretin, the synchronicity of change of hypocretin and estradiol both in hypothalamus and plasma was verified in female rats. With the presence of estradiol, there is specific binding between human ER and the Hypocretin-ERE. Expression of ER combined with estradiol repressed hypocretin promoter activity through the ERE.

**Conclusions:** we defined that estradiol may affect hypocretin neurons in the human hypothalamus via ER binding to the Hypocretin-ERE directly, which may thus lead to the sex-specific pathogenesis of depression.

## Introduction

Depression is a worldwide mental disorder with a series of symptoms affecting cognitive, somatic, affective, and social processes. Previously, our studies reported that both hypocretin-1-expressing neurons and hypocretin-1 plasma were increased in depressed patients compared to controls (Li et al. 2021; Lu et al. 2017). The neuropeptide hypocretin is exclusively synthesized in the hypothalamus, consisting of 2 types: hypocretin-1(the major type in the brain) and hypocretin-2. Hypocretinergic neurons distribute inputs to several brain areas, including the prefrontal cortex (PFC), locus coeruleus, dorsal raphe nucleus, hippocampus, and hypothalamus (Peyron et al. 1998). The hypothalamic-pituitary-adrenal (HPA) axis has been identified by many as the final common pathway for diverse signs and symptoms of depression disorder (Keller et al. 2017; Swaab et al. 2000). Accumulating evidence suggested that the hypocretin system may participate in the modulation of the HPA axis (Kuru et al. 2000; Moreno et al. 2005; NurmioTena-Sempere and Toppari 2010; Russell et al. 2001; Silveyra et al. 2010; Steiner et al. 2013).

Sex difference has been reported to present in various situations. It participates in regulating the stress response, as well as the prevalence of depression. Specifically, women are twice more likely to suffer from stress-related psychiatric disorders than men, taking Major Depressive Disorder (MDD) and post-traumatic stress disorder (PTSD) for example (KeaneMarshall and Taft 2006; Nestler et al. 2002; PearlsteinRosen and Stone 1997; SheikhLeskin and Klein 2002). There is also evidence for sex differences in the hypocretin system, in which the most of studies (both preclinical and clinical) have shown higher hypocretin system expression in females (CataldiLux-Lantos and Libertun 2018; Johren et al. 2001; Lu et al. 2017; Silveyra et al. 2007; Steiner et al. 2013). On the other hand, gonadal steroids, such as estrogen and androgens, are involved in the sex differences of mood disorders. They take effect in a regulatory manner with substrates of the stress response, including the HPA axis (Bao et al. 2006; Seale et al. 2004). In addition, several studies have indicated the participation of the steroid receptor family in the regulation of the activity of stress-related neuropeptide neurons in the hypothalamus (Bao et al. 2006; Dai et al. 2017; Murakami et al. 2011). Recently, some studies have revealed that there was an interactional impact of hypocretin and estrogen in reproduction, sexual behavior, alertness, and the inner biological clock (de Oliveira et al. 2003; Kiezun et al. 2019; Muschamp et al. 2007), which intrigues our great interest towards the topic of the potential role of estrogen in regulating the hypocretin expression in human.

Here we hypothesized that: estrogen acts through its receptors, estrogen receptor α (ERα) and estrogen receptor β (ERβ), which are the nuclear transcriptional factors. Estrogen receptors and hypocretin are co-localized. The estrogen receptor response elements (EREs) that exist in the hypocretin promoter region may directly regulate the gene expression of hypocretin, leading to increased corticotropin-releasing hormone (CRH) activity, which in turn leads to the activation of the HPA axis, thus giving a possible explanation for the sex differences of the prevalence of mood disorders. To verify our hypothesis, we first investigated the potential presence of co-localization of the estrogen receptors (ERs) in hypocretin-immunoreactive neurons in the lateral hypothalamic area (LHA) in the human lateral hypothalamus and defined a clear colocalization of nuclear ERs in hypocretin immunoreactive neuroblastoma SK-N-SH cells. Secondly, we investigated the relationship between E2 and hypocretin in female rats. Plasma estrogen and hypocretin-1 increased in plasma and hypothalamus during the proestrus stage compared with the diestrus stage. In addition, the level of hypocretin-1 in the hypothalamus of the D stage is inversely proportional to the level of hypocretin-1 in the plasma, and the concentration of hormones in the plasma and hypothalamus keeps a synchronization, and the relationship between hypocretin-1 and estrogen is always in direct proportion, that is, hypocretin-1 increases and estrogen also increases, whether detected in the hypothalamus or plasma. After we explored the regulating role of exogenous estrogen on hypocretin gene expression via ERs and subsequently found an activating effect of ERs on human hypocretin expression.

## Material and methods

### Clinical experiment

Seventy-five patients with first-episode drug-naïve MDD (male MDD, n=32, female MDD, n=43) were recruited through outpatient and inpatient visits to the Department of Mental Health of the First Affiliated Hospital, College of Medicine, Zhejiang University. The inclusion and exclusion criteria used were provided in supplementary material. Seventy-four controls (male CTR, n=33; female CTR, n=41) were well-matched with MDD in sex, education, age, and marriage. The controls were recruited from the local community through advertising and subsequently evaluated through the MINI International Neuropsychiatric Interview to ensure that they did not meet any of the DSM-IV psychiatric criteria. Other inclusion and exclusion criteria were the same as those used for the MDD group. All participants signed written informed consent forms. This study was approved by the Ethics Committee of the Department of Mental Health of the First Affiliated Hospital, College of Medicine, Zhejiang University.

### Blood sample collection and plasma hypocretin-1 level measurement

Approximately 5 ml of blood from MDD and CTR were collected and placed in an EDTA tube, followed by centrifugation for plasma collection and storage at -80 ◦C. Plasma hypocretin-1 levels were measured using a commercially available enzyme-linked immunosorbent assay kit (Phoenix Pharmaceuticals Inc., Burlingame, California, USA) according to the manufacturer’s instructions. Plasma was extracted through C-18 columns before testing. The extracted protocols were described in our previous papers (Jin et al. 2020).

### Double labeling of ER and hypocretin in lateral hypothalamic from postmortem tissue

The hypothalamic of a depressive subject was studied. After the autopsy, the hypothalamic was fixed in 4% formaldehyde at room temperature, dehydrated, and embedded in paraffin. After fixing, use a Leitz microtome to cut the paraffin block into sections at a thickness of 6 μm. Then the sections were mounted on SuperFrost/Plus (Menzel, Germany), that, dried for 48 hours on a hot plate at 41 °C and then followed by 24-36 hours in an oven at 37 °C. The sections were deparaffinized in xylene and rehydrated by a series of decreasing ethanol concentrations followed by rinsing in distilled water. Equilibrate the sections in 0.05 M Tris buffer pH 7.6 for 5 mins. Preheat brain sections in the microwave until boiling and then heat the sections in the microwave at a power of 800W for 10 mins. After cooling down, incubate the slides in anti-hypocretin (1:10000, H-003-30, Phoenix Pharmaceuticals, USA) and ERα/ERβ (1:50, NBP2-59201, Novus biological, USA; cSAB2702145, Millpore, USA) diluted in SUMI for 1 hour at room temperature followed by overnight incubation at 4□ in a moist chamber. On the second day, the sections were washed in PBS for 3 x 5 mins, followed by a second incubation with goat anti-mouse (dilution 1:400 in Supermix) for 60 mins. Combinations of secondary antibodies, anti-rabbit Alexa 488 (1:400, srbAF488-1, Proteintech, China) and anti-mouse Alexa 594 (1:400 dilution, SA00013-3, Proteintech, China) were added to the slides for 1 hour and subsequently washed in PBS (3X5 min each). The cells were then counterstained with DAPI (1:200, Life Technologies, Burlington, ON, Canada) for 2 min. Subsequently, the coverslips were mounted on microscope slides (Merck & Co.).

### Double labeling of ER and hypocretin in PC12 and SK-N-SH cell line

The rat pheochromocytoma cell line-PC12 cells were cultured using DMEM (Hyclone, USA) medium supplemented with 10% fetal bovine serum (Hyclone, USA), and 1% Penicillin/Streptomycin. Human neuroblastoma SK-N-SH cells were cultured in DMEM medium supplemented with 20% fetal bovine serum, and 1% Penicillin/Streptomycin. Both cell lines were incubated at 37°C in a 5% CO_2_ incubator. Both cell lines were planted in a 24-well plate with cell slides. The cells were fixed in cold methanol for 15 min, permeabilized with 0.10% Triton X-100 for 25 min, and washed three times with PBS. A blocking solution (1% BSA in PBST) was applied for 30 minutes at 37□. The purified primary antibody against hypocretin (1:400, sc-80263, Santa Cruz Biotechnology, USA) in 1% BSA was incubated overnight at 4°C. Then continued with a second incubation with goat anti-mouse (1:2000, SA00001-1, Proteintech, China) for 60 min. The slides were incubated at room temperature for 10 min using CY5 working solution and subsequently washed in PBS (3X 5 min). Antigen repair solution (pH = 6.0) should be added to the slide. The slide was put in the microwave oven and repaired it once on medium heat for 5 min. Then dropped blocking solution was at room temperature for 30 min to minimize the unspecific antibody binding. Then purified primary antibody against ERα/ERβ (1:400, SC-8005, Santa Cruz Biotechnology, USA;1:400, SC-53494, Santa Cruz Biotechnology, USA) in 1% BSA was incubated overnight at 4□. On the third day, Donkey anti-Mouse (488) secondary antibody was added to the slides and incubated at 37□ for 45min. Wash the slide with TBST for 3X5 min. Then each slide was dripped with DAPI (1:500), incubated at room temperature for 10 min, and washed with TBS for 3X 5 min. Seal the slide with an anti-fluorescence quenching sealing agent and store it at 4 □ in the dark.

### Effect of estrogen on hypocretin-mRNA expression

Estradiol (E2758, Sigma, USA) was dissolved in 100% chloroform for storage. Before use, it was diluted at a final concentration of 10^-5^ M in 0.001% chloroform. Androgen receptor (AR) antagonist ICI 182,780 (V900926, Sigma, USA) was diluted using 100% DMSO and then used at a final concentration of 10^-7^ M in 0.001% DMSO. SK-N-SH cells were grown in 6-well plates for 12 h until attached to the bottom. Once the cells were grown to 70% confluency, AR antagonists were given to the cells for half an hour, followed by the addition of 10^-5^ nM estradiol for 24 h. Control cells were treated with the same content of chloroform and DMSO while absent of AR. Total RNA isolation and cDNA synthesis were performed as described earlier (Lu et al. 2017).

### Real-time quantitative PCR (RT-qPCR) for mRNA expression

The RNA expression levels of hypocretin were detected by using RT-qPCR according to previous protocols (Lu et al. 2015). Briefly, commercial RNA isolation kits (EZBioscience, USA) were used to collect total RNA from the brain tissue. 4× EZscript Reverse Transcription Mix II kit (EZBioscience, USA) was used to synthesize cDNA. RT-qPCR was performed using 2× Universal SYBR Green Fast qPCR Mix (RK21203, ABclonal, China) in a CFX384 Touch Real-Time PCR Detection System (Bio-Rad, USA). All procedures were conducted following the manufacturer’s instructions. The expression levels of individual genes have been normalized to β-actin expression and then calculated by using the 2^−ΔΔCt^ method. All primer pairs were designed with the program Primer Premier 5.0.

### Chromatin immunoprecipitation assay kit (ChIP)

To further explore the activation of ERs by hypocretin, we assessed the binding of ERs to the promoter DNA by ChIP. ChIP assays were performed using a SimpleChIP Enzymatic Chromatin IP kit brought from Cell Signaling Technology (9003, USA). HeLa cells were treated with estradiol (10^-5^M) to induce ERs and cross-linked with 37% formaldehyde for 10 min at room temperature. Immunoprecipitation was performed with anti-ERα/ERβ antibody and anti-hypocretin-1 antibody, and after reverse cross-linking immunoprecipitated DNA was purified by PCR using primers surrounding the EREs sites. Then nuclease digests DNA to get the optimal fragmented chromatin and is subjected to sonication (SONICS-VCX-750, USA). After that, the DNA was fragmented. The fragmented chromatin was around 300-1000 bp as analyzed on agarose gels. ChIP was performed with mouse anti-ERα (NBP2-59197, Novus biological, USA), mouse anti-ER β (cSAB2702145, Millpore, USA), rabbit anti-histone H3 (4620, Cell Signaling Technologies, USA), and normal rabbit IgG (2729, Cell Signaling Technologies, USA) at 4□ overnight. Immunocomplexes were recovered by incubation with blocked protein G beads. After reverse cross-linking and DNA purification, using Power SYBR green with primers for ERs binding sites in the hypocretin promoter (ER), immunoprecipitated DNA was then quantified by real-time PCR. Using the comparative CT method, we calculated fold enrichment based on the threshold cycle (CT) value of the IgG control.

### Electrophoretic mobility shift assay (EMSA)

To determine whether the human hypocretin promoter region contains potential estrogen-responsive elements (EREs), the human hypocretin gene promoter region was searched for consensus ERE motifs (the nucleotide sequences of hypocretin were provided by the National Center for Biotechnology Information database at https://www.ncbi.nlm.nih.gov/gene/3060). At position +1320 bp and +1186bp of the human hypocretin gene promoter, there were two sequences (called site a and site b) of ERα and at position +260bp (site c) of ERβ, respectively in the present study, closely resembling an ERE (site a resemblance rate: 80.8%; site b 205 resemblance rate: 80.3%; site c rate Figure 4A). Probes were designed based on the candidate EREs.

The experimental methods were carried out according to instruments: LightShift™ Chemiluminescent EMSA Kit (20148, Thermofisher, USA). As seen in Figure 3, incubation of 293T nuclear extracts (3ug) with 50nM biotin-tagged probes (Lane 2) was followed by the addition of the same sequences with the non-biotin labeled probe as a competition group (Lane 3). Furthermore, as a control of the competitive reaction, a mutant non-biotin-labeled probe was used (Lane 4). As in the supershift experiments, we also add ERα antibodies to the process (Lane 5). The reaction and detection were carried out according to the methodology used in our previous research (Hu and Chen, 2019). The reaction and detection were carried out according to the methodology used in prior research(Hu et al. 2019). In a nutshell, the reactions were loaded onto a 6% DNA Retardation gel (EC6365BOX, Thermo Fisher, USA) that had been pre-run in 0.5X TBE at 100 V for 1 hour. Then the electrophoretic transfer of binding reaction to a nylon membrane (RPN303B, GE, Amersham, USA) and cross-link DNA to the membrane with a UV transilluminator with a 312 nm lamp for 10 min. The biotin end-labeled DNA was detected using a streptavidin-horseradish peroxidase combination and a light shift chemiluminescent substrate and then exposed membrane to the ChemiDoc Touch Image system (Bio-Rad, Hercules, California) as like western blots did.

### Luciferase assay

To investigate the functional interaction of ERs on the hypocretin, luciferase assays were performed. Four segments of the human hypocretin gene promotor sequence were amplified from genomic DNA to create the hypocretin luciferase reporter plasmids: I a whole segment (from -1434 bp to +76 bp), which contained both potential EREα sites A & B, as well as the potential EREβ site a. ii) a shorter fragment didn’t contain site A (from -1234bp to+76bp). iii) a fragment didn’t contain EREα sites A and site B (from -434bp to +76bp) but only contains EREβ site a, iv) a shorter fragment with no potential ERE sites (from -34bp to +76bp). The schematic representation can be seen in Figure 3E. The fragments were amplified by PCR, and the details and primers were shown in the supplementary materials. The recombinant plasmid was sequenced by Tsingke Biotechnology Co., Ltd. 293T cells were cultured in a 24-well (4×10^5^ cells per well) plate for 24 h before transfection, and then cells were transfected with the fragments which were cloned into the pGL4.10 luciferase reporter basic vector (E1751, Promega, USA), and the PRL-TK reilla luciferase reporter constructs were used as an internal control for plasmid transfection efficiency, pcDNA3.1 plasmid was used as control, together with the plasmids ERα or ERβ, respectively by Lipo8000 (C0533, Beyotime, China). The luciferase activity was then determined using a dual-luciferase assay report kit (E1910, Promega, USA) by a Microplate reader in accordance with the manufacturer’s instructions after transfection for approximately 24 hours.

#### Animals study

Female Sprague–Dawley (SD) rats of approximately 8 weeks of age in this experiment, weighing 220–250 g were housed in a 12h light–dark cycle (lights on at 08:00 h). All the animals were given food and water ad libitum. All the procedures of our animal study were approved by the local animal care committees in accordance with the relevant regulations and laws. At least 3 consecutive regular estrus cycles of 4 or 5 days were monitored by taking daily vaginal smears for the female rats between 09:30 h and 10:00 h. Divide female rats into proestrus (n=7) stage and diestrus (n=6) stage. Female rats were sacrificed by decapitation at the moment that either the proestrus or diestrus was reached. Trunk blood was collected into ice-chilled tubes containing EDTA and then centrifuged at 3000 rpm for 10 min, after which the plasma was separated and stored at −80 □ until hormone measurements were performed. Also, at this time the rat brain was rapidly removed and the hypothalamus dissected. The hypothalami were immediately frozen in liquid nitrogen and then stored at −80□ until next assayed. The dissection and freezing procedure was finished within 3 min after decapitation. For the current study, the enzyme-linked immunosorbent assay method was used (hypocretin-1 kit from Cloud Clone Corp, cEA607Ra, China, and Rat estradiol kits from mlbio, m1002891, China)

#### Statistics analysis

Normal distribution was analyzed by the Shapiro–Wilk test. The Mann–Whitney U test was used to determine nonnormal distribution variables between the two groups, differences among the 3 groups were tested with Kruskal-Wallis (K-W). And t-test and one-way ANOVA were used to determine significant differences in normal distribution variables Receiver operating characteristic curve (ROC) analysis was used to evaluate the diagnostic performance of plasma hypocretin-1 levels. The area under the curve (AUC) is used to assess the ROC effect. All analyses were performed using SPSS (version 22.0, IBM) with a significant P value< 0.05.

#### Ethics approval

The authors assert that all procedures contributing to this work comply with the ethical standards of the relevant national and institutional committees on human experimentation and with the Helsinki Declaration of 1975, as revised in 2008. All subjects signed an informed consent. The study was approved by the Medical Ethics Committee of the First Affiliated Hospital of Zhejiang University School of Medicine (2018-347). And the study was registered in the clinical trial (ChiCTR1900024858).

## Results

### Clinical characteristics of patients

The demographic and clinical characteristics of patients are presented in Table 1, No significant differences were found in sex, age, and education years between in MDD and CTR. Plasma level of hypocretin-1 was significantly increased in the MDD group compared to CTR (27.20pg/ml vs 12.77 pg/ml, P<0.001). Female MDD showed an increase than female CTR (35.40 pg/ml vs 10.27 pg/ml, P<0.0001, Figure 1d) while no significant changes between male MDD and male CTR (16.17 pg/ml vs 16.05 pg/ml, P=0.999). To be noted, ROC analysis revealed an optimal diagnostic value for plasma hypocretin-1 levels in female MDD patients who were drug-naive (AUC:0.766; 95% CI:0.667-0.865). The Yoden’s index was used to determine the optimal cut-off value of hypocretin-1 as 16.755pg/ml, with a sensitivity (0.628) and a specificity of (0.786, Figure 1e).

**Figure 1:**
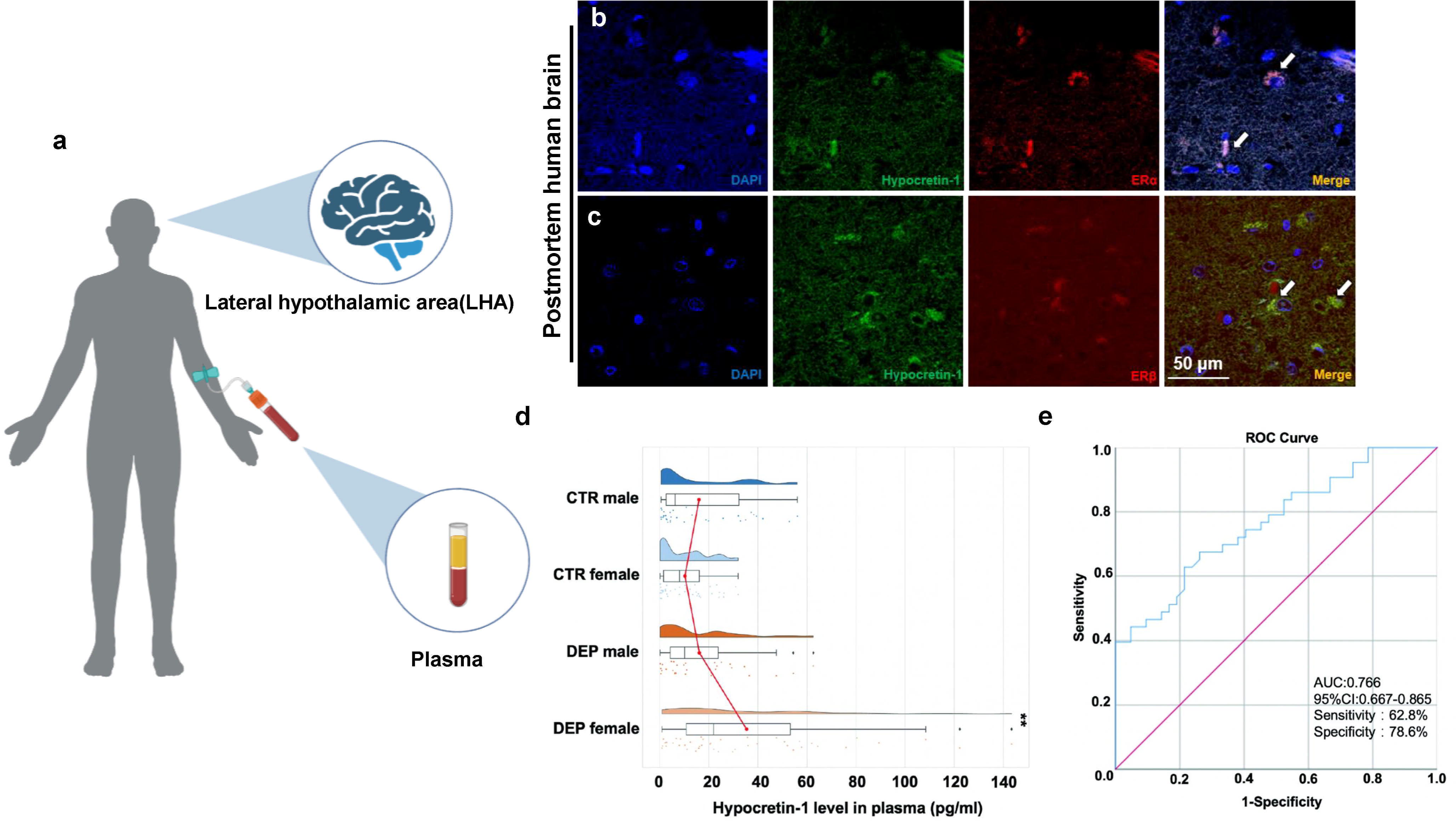
**a.** Show the location of the object of study. **b-c.** Double-labeling immunofluorescence staining showed co-localizations of Estrogen receptor α (ERα) and Estrogen receptor β (ERβ) with hypocretin-1 were shown in a postmortem depressive patients’ lateral hypothalamic area (LHA). **d.** Changes of plasma Hypocretin-1 in female depression (DEP), female control (CTR), male DEP and male CTR. **E.** ROC analysis for plasma hypocretin-1level in female DEP patients.

**Figure 2:**
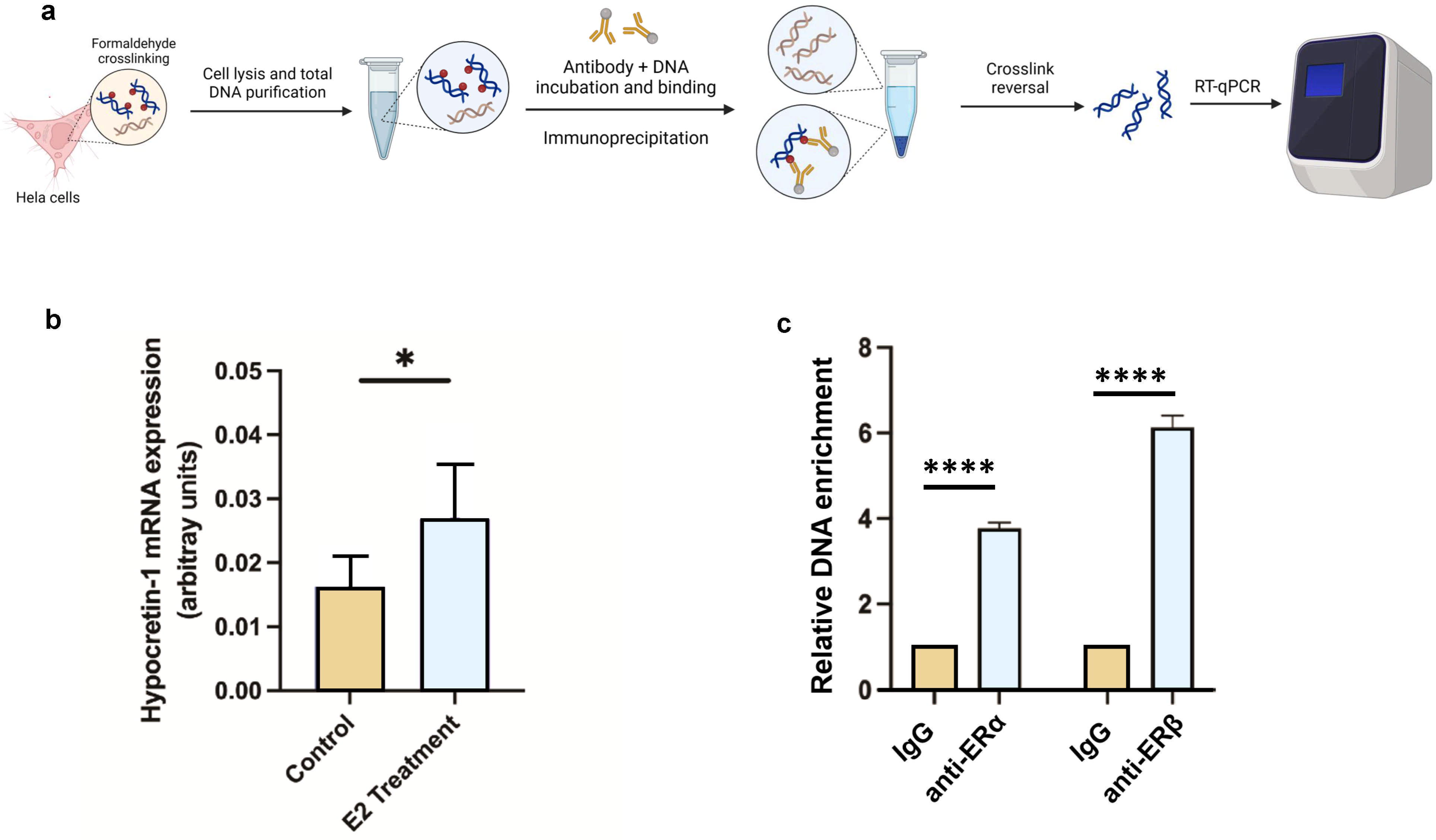
**a.** The experimental model of ChIP. Immunoprecipitated complexes were detected by PCR analysis. Only 293 FT cells co-transfected with POXTA and incubated with anti-ER antibody demonstrated PCR product when Primer A was used. However, cells co-transfected with POXTAm and incubated with anti-ER antibody did not show the PCR products when Primer B was used. In addition, neither untransfected cells treated with anti-AR antibody nor cells co-transfected with POXTA and incubated with IgG demonstrated PCR products when Primer A was used. **b** Comparison of hypocretin-1 mRNA levels in SK-N-SH cells between control group and experiment group with adding the androgen receptor (AR) antagonist ICI 182,780 to the medium, experiment group following adding 10-5 nM estradiol for 24 h while control group adding same content of DMSO solvent to the medium. Data are presented as mean +/- SEM from three independent experiments. * p =0.05. E2 treatment: estradiol treatment. **c**. ChIP assay showing binding of ERs to the promoter. Precipitated DNA was assayed by qPCR with specific primers for amplification of EREs of the hypocretin promoter.

**Table 1:** Demographic and clinical characteristics.

| Measure | CTR-male | CTR-female | MDD-male | MDD-female | t/$\chi^2$/F | Adjusted P value |
| --- | --- | --- | --- | --- | --- | --- |
| Age (year) | 25.91±4.795 | 28.81±7.970 | 26.41±5.988 | 27.16±5.723 | 1.812 | 0.612 |
| Number | 32 | 42 | 32 | 43 |  |  |
| Education years | 15.03±1.257 | 15.07±1.583 | 14.72±2.036 | 14.49±1.980 | 0.930 | 0.818 |
| Course of disease<br>(month) | | | 0.6303±0.6107 | 0.6416±0.7044 | -0.073 | 0.942 |
| HAMD | 2.156±2.034 | 2.095±2.377 | 23.81±4.410<sup>****</sup> | 24.42±4.625<sup>####</sup> | 111.9 | <0.0001 |
| HAMA | 1.875±2.673 | 1.167±1.666 | 21.39±5.990<sup>****</sup> | 12.17±5.357<sup>####</sup> | 112.8 | <0.0001 |

a.t-test; b. $\chi^2$ test; \* comparison between male CTR with male MDD, # comparison between female CTR
with female MDD. \*\*\*\*, P<0.0001, ####, P<0.0001. MDD: major depressive depression.

### Co-localization of ER and hypocretin-1 in neurons of the LHA in postmortem, SK-N-SH cells, and PC12 cells

Frontal section of the ateral hypothalamus area (LHA) in subject (#00182) was stained for hypocretin and ERα (Figure 1b) or ERβ (Figure 1c). hypocretin-1 was present both in the cytoplasm and fibers, while ERα and ERβ were also found in the cytoplasm and fibers of hypocretin-expressing neurons. In addition, We also investigated the co-location in neurocyte cell lines, hypocretin-1 showed clear co-localizations of endogenously expressed ERα and ERβ in PC12 cells (Supplemen Figure 1a-b) and SK-N-SH cells (Supplement Figure 1c-d).

### Estradiol increases hypocretin-mRNA in vitro

After pre-incubation of the SK-N-SH cells with the AR antagonist followed by estradiol, significantly elevated hypocretin-mRNA levels were observed (student’s t-test, p =0.032, Figure 3b).

**Figure 3:**
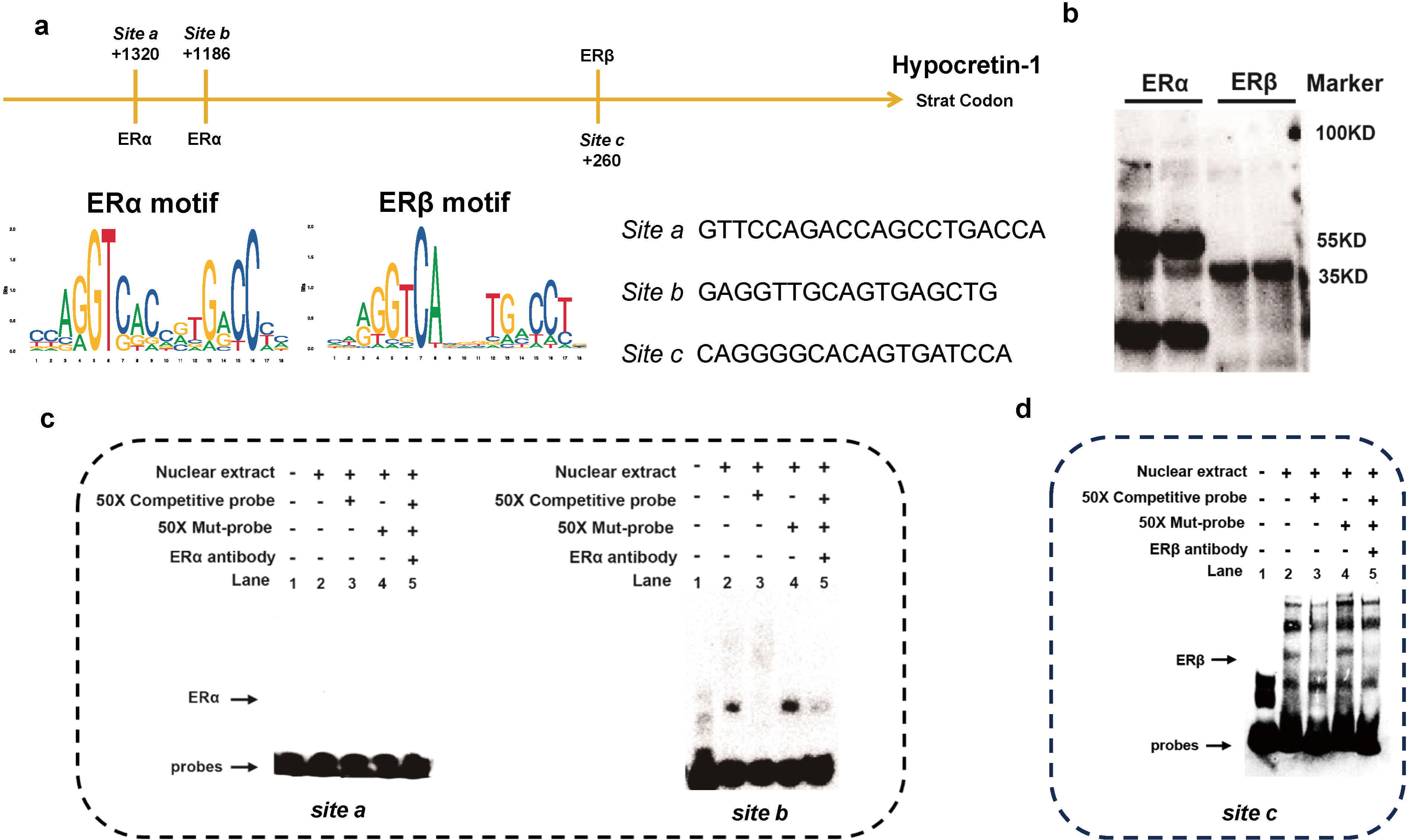
**a.** The schematic shows the promoter of the human hypocretin gene, as well as potential ERE binding site, including two ERE-α binding sites at +1320 (site a) and +1186 (site b) and an ERE-β binding site was at +260 (site c). The site positions were relative to the start codon. **b.** ER-α displayed two variations at roughly 35 KD and 70 KD. c**-d.** Electrophoretic mobility shift assay (EMSA) of predicted estrogen responsive elements (EREs). ERE-potential α sites (site a&b) and β site (site c) incubated with labeled human hypocretin. Site b (line 2) strongly combines with nuclear ER-α extracts and site c (line 2) combines with nuclear ER-β extracts from transfected 293T cell lines, as indicated by the arrow. The band vanished when 50X competition probes (same sequences as the intended probe but not biotin-labeled) were added to the combination reaction mix (site b, lane 3; site c, lane 3). When we replaced the competition probes with Mut-probes (mutated sequences with no biotin label), the band reappeared (site b, lane 4; site c, lane 4). When antibodies anti-ERα or anti-ERβ were added to the combination mix, the band shrank even though there was no supershift band (site b, lane 5; site c, lane 6).

### Identification of ERE-α and ERE-β binding sites in human hypocretin-1 promoter

The protein-oligonucleotides complexes were detected by EMSA assay, and the results showed that only site b (probe 2) and site c (probe 3) but not site a (probe 1) of hypocretin promoter strongly bound to the nuclear extract from ERα or ERβ overexpressing 293T cells, respectively, and the ERα-hypocretin promoter complexes and ERβ-hypocretin promoter complexes binding bands shown as the arrow pointed (see Figure 4c, site b, lane 2; Figure 4d, site c, lane 2; respectively). The binding bands vanished when the complex was incubated with a 50 excess of unlabeled probe 2 or unlabeled probe 3, respectively (see Figure 4c, Site b, lane 3; Figure 4d, site c, lane 3) and reappeared when the unlabeled probes were mutated (see Figure 4c, Site b, lane 4; Figure 4d, site c, lane 4). The specific bands shrank even though there was no obvious super-shifted band after the addition of the antibody of ERα or ERβ (see Figure 4c, site b, lane 5; Figure 4d, site c, lane 5). Herein, EMSA results strongly indicated that ERα and ERβ physically bind to ERE-α site b and ERE-β site c on the hypocretin promoter, respectively.

**Figure 4:**
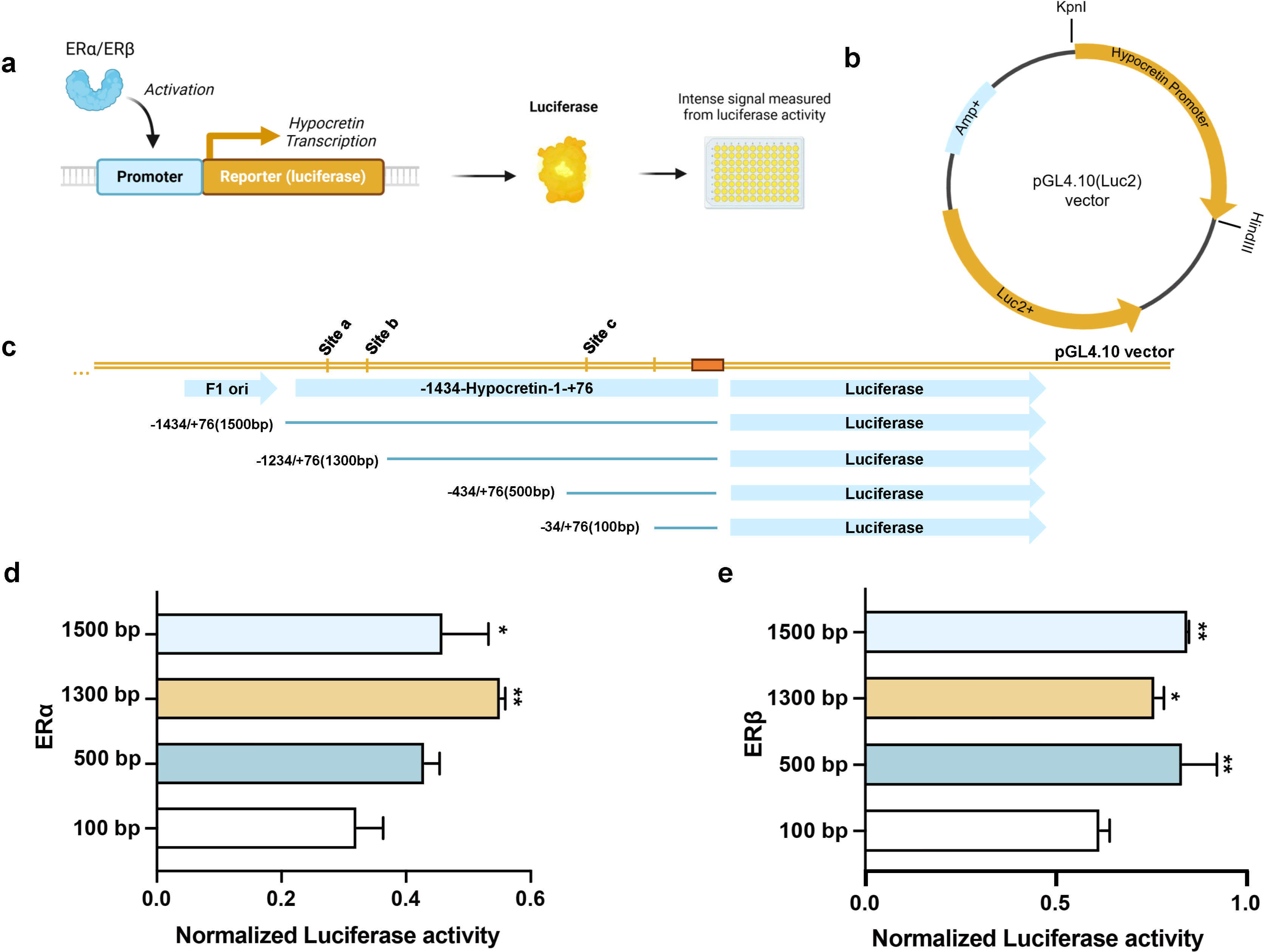
**a.** The experimental model of luciferase. **b.** The vector and double cleavage site. **c.** To construct the hypocretin luciferase reporter plasmids, four different sites fragments from the human hypocretin gene promotor sequence were amplified from genomic DNA by PCR. **d.** Compared to 100bp pGL 4.10, hypocretin transcriptional luciferase activity presents a significant increase in cells transfected with ERα plasmids together with 1500bp pGL 4.10 (*p* = 0.001) and 1300bp pGL 4.10 (*p* = 0.029), but not in cells transfected with 500bp pGL 4.10. **e**. Compared to 100bp pGL 4.10, hypocretin transcriptional luciferase activity presents a significant increase in cells transfected with ERβ plasmids together with 1500 bp pGL 4.10 (*p* = 0.003) and 1300 bp pGL 4.10 (*p* = 0.038) and 500 bp pGL 4.10 (*p* = 0.002). (\**p*<0.01, \*\**p*<0.001)

Investigation of evidence of direct binding of ERs to the hypocretin promoter by ChIP assay. Which shows binding of ERs to the promoter. HeLa endogenous ERs was induced by estrogen treatment for 48 h and then cross-linked. ChIP of fragmented DNA was performed with ER-antibody, and ER-antibody or control IgG. Precipitated DNA was assayed by qPCR and showed ERE binds to hypocretin promoter (all p<0.0001, Figure 3c).

### ER upregulates the hypocretin promoter

Hypocretin transcriptional luciferase activity presents a significant increase in cells transfected with ERα plasmids together with -1434bp to +76bp pGL 4.10 (p = 0.001) and -1234bp to +76bp pGL 4.10 (p = 0.029), but not in cells transfected with - 434bp to +76bp pGL 4.10 compared to -34bp to +76bp pGL 4.10 (Figure 5d). In addition, hypocretin transcriptional luciferase activity presents a significant increase in cells transfected with ERβ plasmids together with -1434bp to +76bp pGL 4.10 (p = 0.003) and -1234bp to +76bp pGL 4.10 (p = 0.038) and -434bp to +76bp pGL 4.10(p = 0.002) compared to -34bp to +76bp pGL 4.10 (Figure 5e).

**Figure 5:**
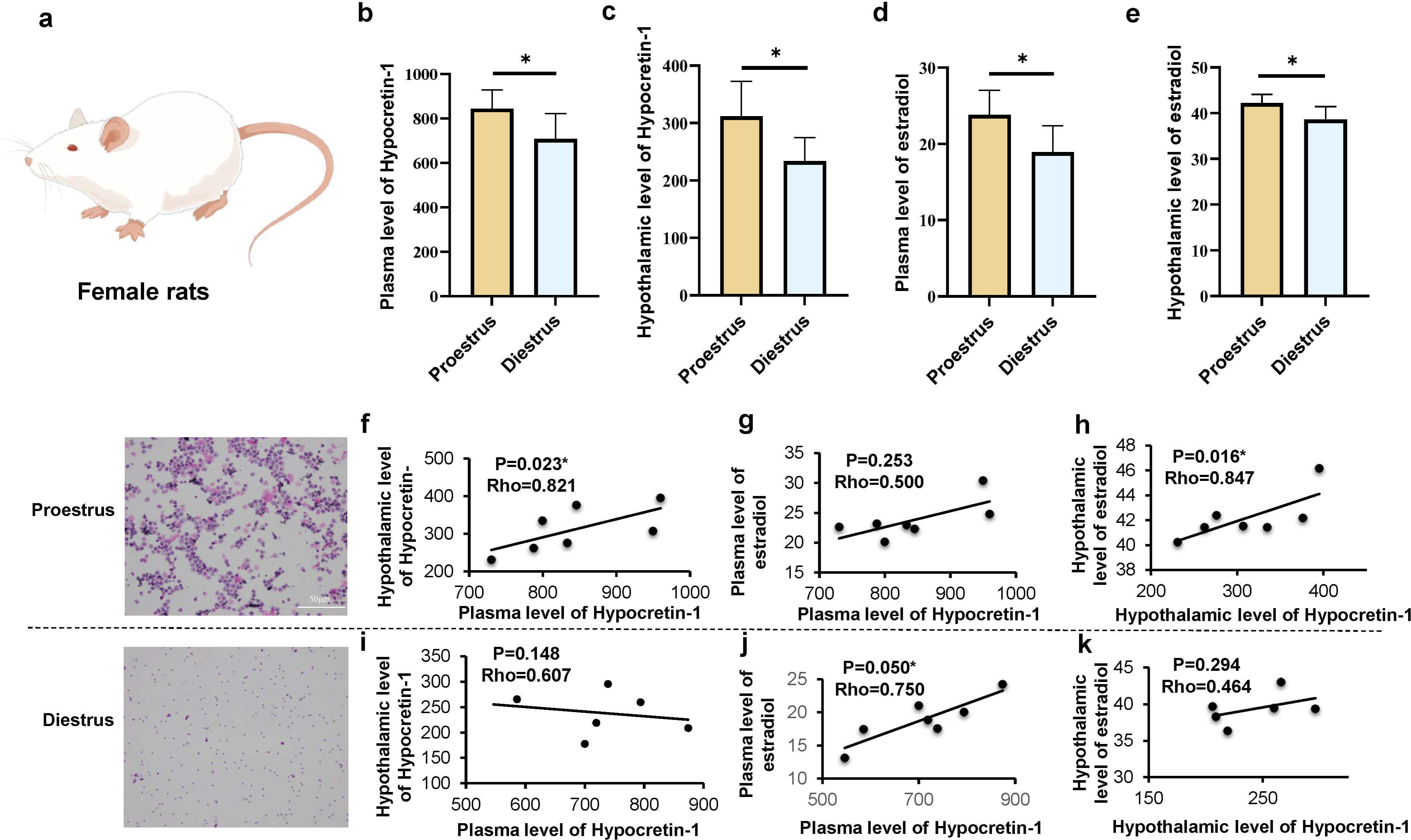
Levels of hypocretin-1 and estradiol in rat plasma and hypothalamus. **a.** An experimental female rat was shown. **b.** Changes in plasma hypocretin-1 of female rats in proestrus (P) and diestrus (D) stage, significantly increased at P stage. **c.** Changes in hypothalamic hypocretin-1 of female rats in P and D stage, significantly increased at P stage. **d.** Changes in plasma estradiol (E2) of female rats in P and D stage, significantly increased at P stage. **e**. Changes in hypothalamic estradiol of female rats in P and D stage, significantly increased at P stage. **f-h**. Corrlations of hypothalamic, plasma hypocretin or estradiol in the proestrus (P) stage. **i-k.** Corrlations of hypothalamic, plasma hypocretin or estradiol in the diestrus (D) stage.. (* *P*<0.05).

### Plasma and hypothalamic level of hypocretin-1 and estradiol in proestrus and diestrus

In female rats, the plasma level of hypocretin-1 and estradiol was increased in prxoestrus than in diestrus (p = 0.039, p = 0.018, respectively, Figure 6b, 6d), Hypothalamic level of hypocretin-1 and estradiol also higher in proestrus compared to diestrus (p = 0.017 and p = 0.016, respectively, Figure 6c, 6e). In proestrus, we found a positively correlation between hypothalamic hypocretin-1 and plasma level of hypocretin-1(p = 0.023, rho = 0.821, Figure 6f) and positively correlation between hypothalamic E2 and hypothalamic hypocretin-1(p = 0.016, rho = 0.847, Figure 6h). In diestrus stage, Only a positive correlation between the plasma level of estradiol and the plasma level of hypocretin-1 was found (p = 0.050, rho = 0.750, Figure 6j).

## Discussion

Our previous series of postmortem material so far confirms the presence of more hypocretin expression in the hypothalamus in female depressive patients(Lu et al. 2017). Suggesting estrogen might regulate the human hypocretin system. A series of research methods and research objects to explore its possible mechanism. We confirmed these changes in clinical female drug naïve-depressive patients, which showed higher plasma hypocretin-1 in female depression. For the first time, we found the double-labeling of ERα and ERβ with hypocretin-1 in the human lateral hypothalamus, Moreover, colocalization in the neuroblastoma SK-N-SH cell line and PC12 cell line, which was reported to have endogenous ER and hypocretin expression (Li and Qiu 2015; Wieland et al. 2002), we observed now that estradiol-induced could increase hypocretin-mRNA expression through ERs. Notable, it is not necessary to show the correlations between ER expression in depression disorder, since the expression of hypocretin could be regulated by multiple factors, such as ER or transmitters. Then it was also observed that the hypothalamic levels of estradiol and hypocretin in female rats during the proestrus stage were positively correlated. Following these observations, we concentrated on two candidate sites and one in the present study, which all are located upstream of the transcriptional start site ATG. the two functional ERE binding sites in the hypocretin promoter were verified by ChIP, EMSA, and luciferase. Here, we identified ERE sites on the human hypocretin promotor. Though no previous evidence has directly indicated this relationship, studies have shown that the hypocretin system is controlled by estrogens, and the expression of components of the hypocretin system is also regulated in a sex-dependent manner. In adult gonadectomized male rats, an increased expression of hypocretin-1 receptors in the pituitary and testosterone replacement could reverse this effect. On the other hand, no effect of gonadectomy and steroid replacement could be found on the expression of components of the hypocretin system in the hypothalamus of male and female rats (Johren et al. 2003) although hypothalamic expression of prepro-hypocretin and hypocretin receptor is higher in female rats compared to males (Johren et al. 2002).

Hypocretin contributes to the regulation of various physiological functions, including sleep and wakefulness, reward-related behavior, arousal, energy homeostasis, as well as neuroendocrine function(Milbank and Lopez 2019). Pu et al hypocretin could stimulate or inhibit pituitary luteinizing hormone (LH) secretion dependent on the presence of ovarian steroid hormones(Pu et al. 1998), In addition, a sexually dimorphic expression of hypocretin/prepro-hypocretin and hypocretin receptors in the hypothalamus was found (Lu et al. 2017). In accordance with our previous results (Lu et al. 2017), hypocretin-1 levels were reported to be higher in the lateral and posterior hypothalamus of female rats compared to male rats (Taheri et al. 1999). Hypothalamic prepro-hypocretin mRNA levels were also higher in female rats compared to males (Johren et al. 2002). Moreover, the expression of the hypocretin-1 receptor was observed to be higher in the hypothalamus of female rats compared to male rats (Johren et al. 2001). Our previous research showed that postmortem results indicated that women with depression had significantly more hypocretin neurons positive than healthy controls, while there was no significant change in men(Lu et al. 2017).

It is worth noting that there have been many reports that hypocretin regulates sex hormones, but little is known about the reverse regulation. The studies on humans and animals have found that acute changes in estrogen can lead to an overactive hypocretin system, and cause depression and anxiety symptoms(Federici et al. 2016; Sundström Poromaa et al. 2017). Under stress, estrogen will affect the balance of the HPA axis, making female HPA axis activity more easily hyperactive. In clinical, plasma hypocretin-1 levels were significantly higher in postmenopausal women after oral estrogen than in the control group(Cintron et al. 2017). In the preclinical studies, It has found after injecting hypocretin-1 into the lateral ventricles of normal groups and ovariectomized rats, luteinizing hormone secretion increased in normal rats, while significantly decreased in ovariectomized rats(FurutaFunabashi and Kimura 2002). Hypocretin has been shown to have a stimulatory effect on release of gonadotropin-releasing hormone (GnRH) from rat hypothalamic explants in vitro, mediated via the HCRTR1, coupled with activation of the PKC-, MAPK- and PKA-signaling pathways(Sasson et al. 2006).

Several studies have demonstrated that hypocretin are involved in stress regulation of depression through the hypothalamic-pituitary-adrenal (HPA) axis(Pu et al. 2022). Intraventricular injection of hypocretin-1 can activate the expression of CRH-mRNA in the PVN and increase the plasma Adrenocorticotropic Hormone (ACTH) and CORT levels(Kuru et al. 2000). The sex difference in the alterations of hypocretin, which happens in the framework of the stress hypothesis, was further supported by our animal study, showing a significant positive correlation between hypothalamic prepro-hypocretin-mRNA and CRH-mRNA only in female CUMS rats(Lu et al. 2017). After treated hypocretin receptor 1 antagonist (SB334867), the HPA axis was significantly inhibited and the occurrence of cognitive impairment was restored in female rats(Grafe et al. 2017).

In postmortem case studies, we had only a single case with co-localization. It should also be noted that the effect of estrogen on hypocretin expression in SK-N-SH cells data are based upon mRNA measurements and have yet to be confirmed on the protein level.

To be noted, our previous studies showed hypocretinergic system showed sex differences in psychiatric disorders, such as depression(Lu et al. 2017) and schizophrenia(Lu et al. 2021). And the potential role of sex hormones such as androgens on the activity of peptidergic neurons other than hypocretin such as CRH neurons, and oxytocin-containing neurons(SwaabBao and Lucassen 2005), indicating that the onset of mood disorders has a great relationship with sex hormones. Hypocretin may be important in the etiology of stress-related psychiatric disorders that present differently in men and women. Thus, targeting hypocretin could potentially ameliorate many phenotypes of stress-related illness in a sex-specific way(Grafe and Bhatnagar 2020).

In conclusion, our results indicated that estrogen may thus participate in regulating hypocretin levels via its receptors, ERα and ERβ, both in health or disease. We conclude that estrogen may directly affect hypocretin neurons in the human hypothalamus via ER binding to the hypocretin-ARE, which may have consequences for the sex-specific pathogenesis of mood disorders.

## Supporting information

STable 1

## Acknowledgments

The figure of the article was drawn using BioRender (https://biorender.com/). The authors are grateful to the Netherlands Brain Bank (Director Dr. Inge Huitinga) for providing human brain material and clinical details. We sincerely thank the support of funds from the National Natural Science Foundation of China (82271561) and the Medical Science and Technology Project of Zhejiang Province (2022RC024) and the Natural Science Foundation of Zhejiang Province (LQ20H090010).

