## Supplementary material for "A direct estrogenic involvement in the expression of human hypocretin": STable 1

**The inclusion and exclusion criteria**

(1) a 17-item Hamilton Depression Scale (HAMD-17) score≥17; (2) 18–45 years old; (3) first onset and not taking any psychiatric drugs before the trial; (4) Han ethnicity; (5) patients who met the Diagnostic and Statistical Manual of Mental Disorders 5 (DSM-5) criteria for current unipolar MDD. The exclusion criteria were as follows: (1) any neurological disease, including organic disease and mental illness other than depression; (2) serious diseases of the kidney, liver, cardiovascular, or respiratory systems; (3) any serious medical condition with major medical intervention anticipated during the experiment; (4) pregnancy or breastfeeding; (5) use of any supplement with potential antidepressant effects; and (6) a serious risk of suicide or violence

**Supplementary Figure 1**

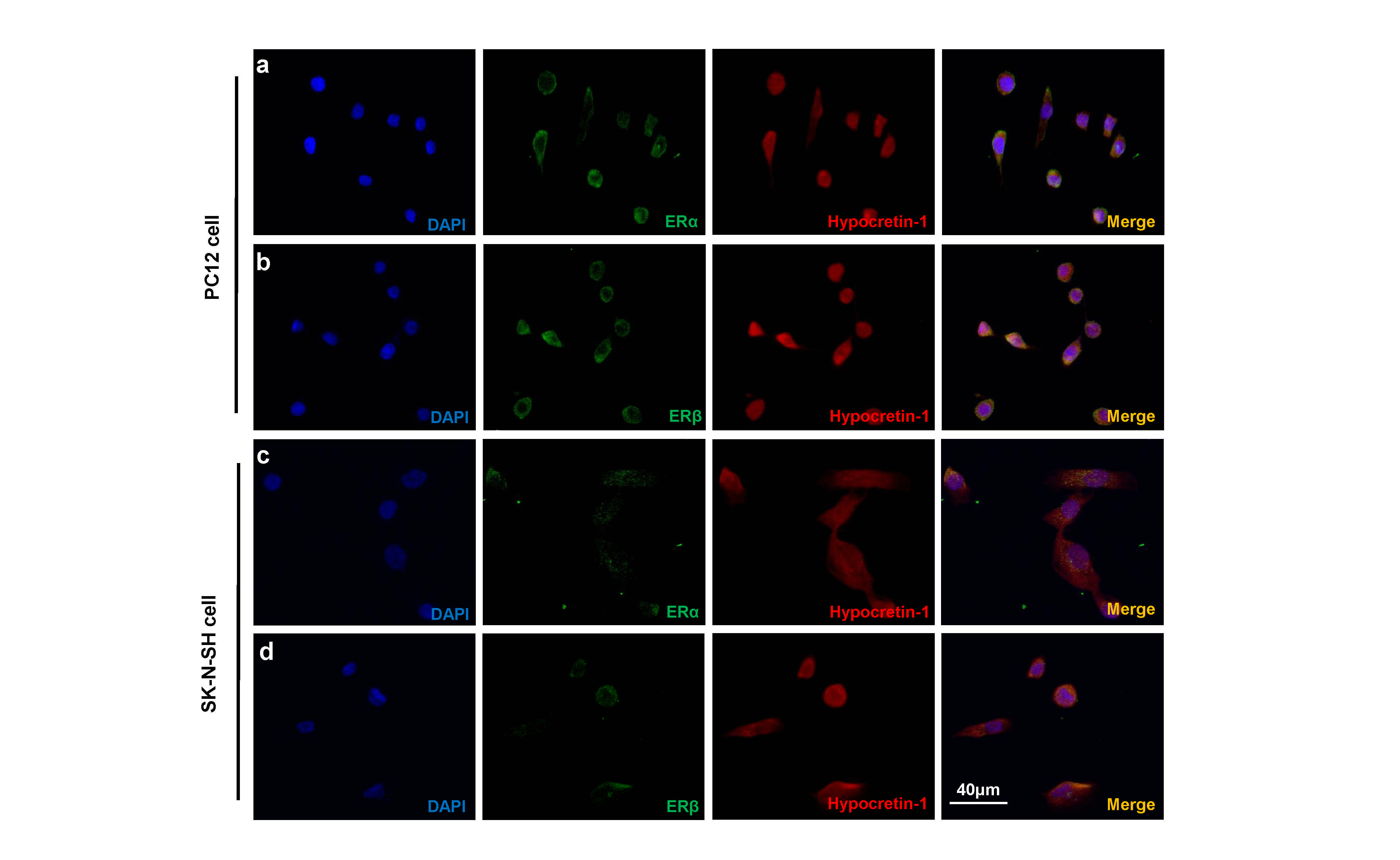

**a-b.** Double-labelling immunofluorescence staining showing co-localizations of ERα and ERβ with hypocretin-1 were shown in PC12 cells, respectively. **c-d.** Co-localizations of ERα and ERβ with hypocretin-1 were shown in SK-N-SH cells, respectively.

**Supplementary Table 1**

Table 1: Demographic and clinical characteristics

a.t-test; b. χ2 test; ^*^ comparison between male CTR with male MDD, ^#^ comparison between female CTR with female MDD. ^****^, P<0.0001, ^####^ , P<0.0001. MDD: major depressive depression.

| Measure | CTR-male | CTR-female | MDD-male | MDD-female | t/χ2/F | Adjusted P value |
| --- | --- | --- | --- | --- | --- | --- |
| Age (year) | 25.91±4.795 | 28.81±7.970 | 26.41±5.988 | 27.16±5.723 | 1.812 | 0.612 |
| Number | 32 | 42 | 32 | 43 |  |  |
| Education years | 15.03±1.257 | 15.07±1.583 | 14.72±2.036 | 14.49±1.980 | 0.930 | 0.818 |
| Course of disease (month) |  |  | 0.6303±0.6107 | 0.6416±0.7044 | -0.073 | 0.942 |
| HAMD | 2.156±2.034 | 2.095±2.377 | 23.81±4.410^****^ | 24.42±4.625^####^ | 111.9 | <0.0001 |
| HAMA | 1.875±2.673 | 1.167±1.666 | 21.39±5.990^****^ | 12.17±5.357^####^ | 112.8 | <0.0001 |
